# Biodiversity-ecosystem function relationships change in sign and magnitude across the Hill diversity spectrum

**DOI:** 10.1101/2022.09.30.510130

**Authors:** Michael Roswell, Tina Harrison, Mark A. Genung

## Abstract

**Summary:** Motivated by accelerating anthropogenic extinctions, decades of biodiversity-ecosystem function (BEF) experiments show that ecosystem function declines with species loss from local communities. Yet, at the local scale, changes in species’ total and relative abundances are more common than species loss. The consensus best biodiversity measures are Hill numbers, which use a scaling parameter, ℓ, to emphasize rarer versus more common species. Shifting that emphasis captures distinct, function-relevant biodiversity gradients beyond species richness. Here, we hypothesized that Hill numbers that emphasize rare species more than richness does may distinguish large, complex, and presumably higher-functioning assemblages from smaller and simpler ones. In this study, we tested which values of ℓ produce the strongest BEF relationships in community datasets of ecosystem functions provided by wild, free-living organisms. We found that ℓ values that emphasized rare species more than richness does most often correlated most strongly with ecosystem functions. As emphasis shifted to more common species, BEF correlations were often weak and/or negative. We argue that unconventional Hill diversities that shift emphasis towards rarer species may be useful for describing biodiversity change, and that employing a wide spectrum of Hill numbers can clarify mechanisms underlying BEF relationships.

## Introduction

A central question in community ecology is, “how will ongoing shifts in biodiversity affect ecosystem function?” In experiments that vary species richness while controlling other community properties, the answer has been clear for some time: ecosystem function has a positive, saturating relationship with species richness [1–3]. There is ongoing interest in “scaling up” research to resolve whether similar patterns hold in natural ecosystems [4,5]. However, richness is not a robust measure of biodiversity in observational data taken from natural ecosystems [6], in large part because most species are rare [7] and likely to be absent from samples. Further, richness often tracks real-world biodiversity gradients poorly, because species composition and abundance can change dramatically with little to no change in observed species richness [8–10]. Therefore, other metrics of biodiversity may provide improved clarity about the real-world linkages between biodiversity and ecosystem function (BEF).

There are both historical and conceptual reasons that BEF research has focused on richness as a measure of biodiversity. Since at least the 1960s, there has been extensive research on how productivity affects species richness [1]. Motivated by intensifying biodiversity loss in the 1980s, declines in richness were (at least implicitly) the global change pattern that seminal BEF studies, with their focus on species loss (e.g., [11]), aimed to understand. This prompted a wave of experiments on how species richness affects ecosystem function [2]. Thus, richness was a natural choice, both because of ecology’s long focus on how richness might *respond* to ecosystem functions like productivity, and because of a collective sense that species loss was the correct, or at least most convenient, way to frame anthropogenic changes in biodiversity. Furthermore, richness was considered a good proxy for functional diversity and redundancy, which were considered the key mechanisms through which biodiversity maintains ecosystem function [12–14]. However, the choice of richness may not have been based on theoretical expectation that richness, rather than other abundance-weighted diversity measures, best described functionally important biodiversity gradients.

Using species richness as the key biodiversity measure poses methodological problems for BEF research, especially when community properties other than richness vary, as in naturalistic systems. Species richness is not only sensitive to the extent and depth of sampling, but also to the distribution of relative abundances in the sampled assemblage. To illustrate this, consider the difficulty of accurately measuring species richness in a community with one hyper-dominant species and many very rare species, versus measuring richness in a community in which abundance is evenly distributed. Richness, like other diversity measures, summarizes the distribution of relative abundances in an assemblage, and when estimated from data, cannot be independent from that distribution, even if such a measure were desirable [6,15]. However, different diversity measures vary in the extent to which they emphasize rare vs. common species, with species richness heavily emphasizing rare species. A unified family of diversity measures, known as “Hill numbers” or “Hill diversities,” summarizes a distribution of relative abundances as the abundance-weighted, generalized mean rarity [16–18]. Hill numbers are governed by a scaling parameter, ℓ, that scales species rarity when computing the mean, and higher values of ℓ afford more leverage, or emphasis, to rare species, while lower values emphasize common species more [18].

The Hill diversity of an assemblage is not a single value, but rather a spectrum that varies continuously across ℓ [6,16] (Figure 1), raising the question of how ecosystem function relates to biodiversity measures with different emphasis on common vs. rare species. While several recent studies have compared whether richness (ℓ = 1), exponentiated Shannon (ℓ = 0), or inverse Simpson (ℓ = −1) best explains ecosystem function [19–22], there has been no examination of how Hill numbers relate to ecosystem function across a wide range of ℓ values. This is a striking knowledge gap because, although nearly all studies of the relationship between biodiversity and ecosystem function have used species richness as a measure of diversity [3], other diversity measures could both better describe real world variation in biodiversity, and also have stronger links to ecosystem functioning. Furthermore, Hill diversities with ℓ > 1, which emphasize rare species even more than richness does, have scarcely been studied at all, not to mention in relationship to ecosystem function. Thus, biodiversity-function studies may be *underestimating* the importance of biodiversity for function by not considering Hill diversities with different emphases on rare and common species via different values of the scaling parameter ℓ.

**Figure 1.**
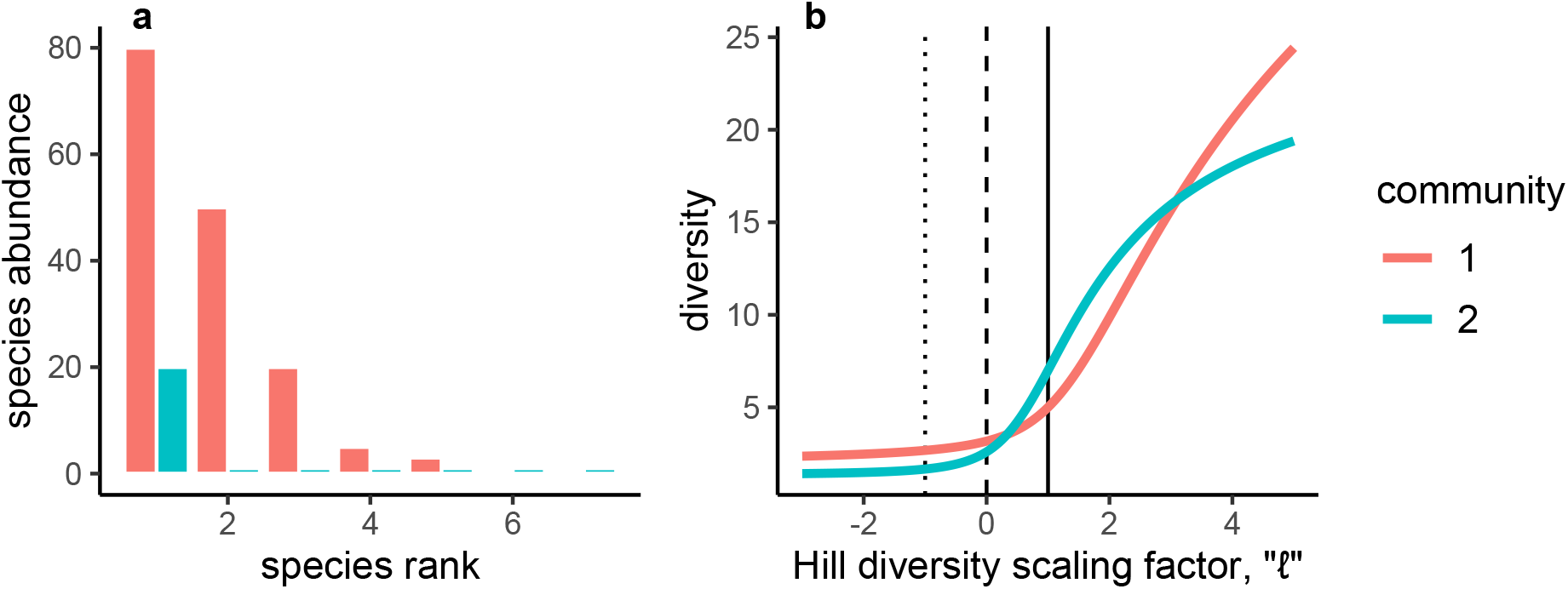
Two hypothetical communities with (a) different species abundance distributions have (b) different diversity profiles. At large negative ℓ values, each diversity profile converges on the inverse proportional abundance of the one most abundant species in the assemblage (inverse dominance). As ℓ values are more positive, each diversity profile converges on the inverse proportional abundance of the one least abundant species in the assemblage (equal to total abundance when the least abundant species is a singleton). Because singletons are ubiquitous in observational data, sample Hill diversities converge on observed abundance with increasingly large, positive values of ℓ. In the example, the red community is more even and more abundant, so its diversity is higher compared to the blue community at both ends of the diversity spectrum. However, the blue community has more species, and therefore is more diverse around richness (ℓ = 1, solid vertical). Other commonly used diversities are inverse Simpson (dotted, ℓ = −1) and exponentiated Shannon (dashed, ℓ = 0). In the diversity literature, it has been less common to explore the right side of this spectrum (i.e., ℓ > 1)

Despite clear declines in richness at the global scale, local changes in biodiversity and their connection to function are likely better captured by measuring total abundance and species’ relative abundances [8,23], for at least three reasons. First, as already discussed, observed richness is a poor predictor of true richness [24], and good estimators of true richness based only on species frequencies in samples may never exist [25]. Thus, even if underlying variation in species richness correlates strongly with, or even drives, ecosystem function, estimating richness from samples could severely obscure the underlying pattern. Second, although observed richness does increase with observed abundance, to the extent that abundance *per se* drives function, Hill diversities that better reflect abundance (i.e., when ℓ >>1) should be stronger correlates of function than richness (ℓ = 1) is. Third, if diversity effects on function are mediated by positive species interactions [3,26–28], more probable and stronger between equally abundant species [29,30], Hill diversities that better reflect the probability of interspecific encounter (e.g. Hill-Simpson diversity, at ℓ = −1 [31]) might explain function better than richness does.

Here, we ask how biodiversity-ecosystem function correlations in observational datasets change in sign and magnitude across a wide range of values of the scaling parameter ℓ. In natural communities, classic BEF mechanisms such as selection and complementarity co-occur with other sources of variation in function, such as abundance, evenness, demography, and environment [32,33].There is no way to standardize BEF modeling approaches across heterogeneous natural systems [34–36], so researchers variously use mathematical partitions [37] or regression and path analysis [32,38] to partially account for subsets of these factors depending on their system knowledge, assumptions, and preferred study focus. In order to focus attention on the possibilities of a wide diversity spectrum, we keep analyses simple and general by presenting the overall correlation between total function and diversity across natural communities, which represents the net effects of many factors. We focus on ecosystem functions that can be expressed as the product of mean per-capita function and total abundance, which works well for many functions [39,40]. As a first step towards a more granular approach, we also present separate correlations between diversity and total abundance [41] and between diversity and per-capita function, which captures selection effects due to shifts in community composition, together with complementarity effects on individual-level function. We explore whether diversity measures other than richness can better explain BEF patterns and potentially help identify BEF mechanisms in natural systems.

In this study, we use observational community datasets on three ecosystem functions to ask:

1. Which values of the Hill diversity scaling factor ℓ produce the strongest biodiversity-ecosystem function correlations?
2. How do biodiversity-ecosystem function correlations change in sign and strength over a wide range of Hill diversities?
3. What is the role of absolute abundance in shaping BEF correlations over the Hill diversity spectrum?

## Materials and Methods

To find how biodiversity-ecosystem function correlations change with different diversity scaling, we used previously published community datasets that recorded both the abundance and function of species at multiple sites. We chose datasets of disparate ecosystem functions and spatial scales: pollination by wild bee visitors to a landscape-scale array of three plant species [42,43], reef fish biomass from dive surveys replicated within 32 globally distributed geographic regions [44,45], and above-ground tree biomass in census plots replicated within four tropical forests [46,47] (Table 1). In each system, total function of a community can be estimated as the summed contribution across species (or individuals) present in the community. Pollination was measured as the product of first, the typical number of pollen grains deposited during a single visit of a focal bee taxon to the focal plant species, and second, the number of observed visits to the focal plant species by the focal bee taxon. Reef fish biomass was measured by visually estimating individual fish body lengths during dive surveys, which were then used to calculate biomass using species-specific allometric equations. Tropical tree biomass was measured by converting observed individual diameter at breast height into biomass estimates using taxon-specific allometric equations that included information about wood density [48,49]. In total, we used 39 community datasets, each consisting of one function measured across a collection of assemblages.

**Table 1.**
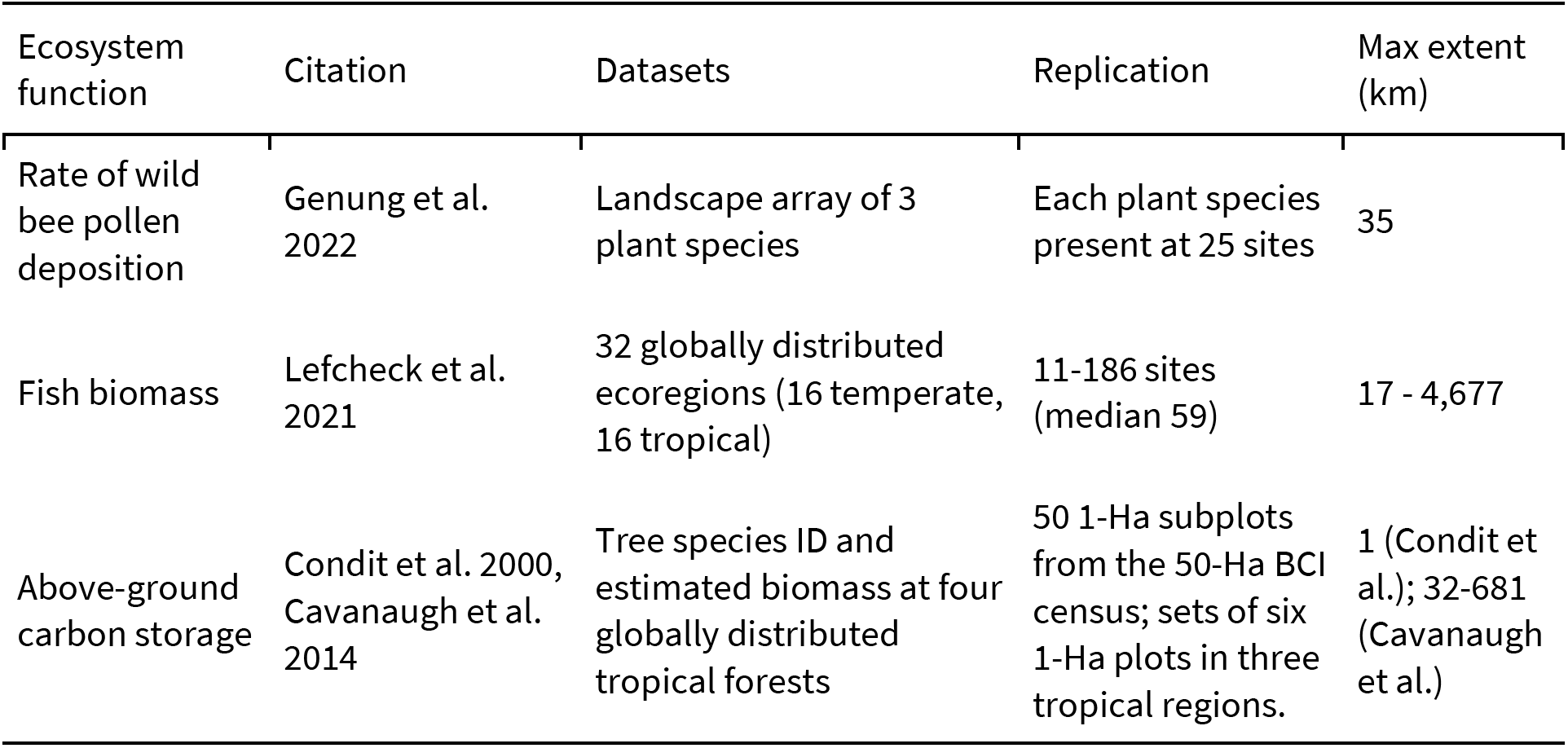
To learn how biodiversity-ecosystem function correlations are affected by different Hill diversity scaling factors, we gathered published, observational community datasets on three ecosystem functions. These were subdivided into a total of 39 community datasets, each including observations of species’ identities, abundances, and functions across replicated sites.

| Ecosystem function | Citation | Datasets | Replication | Max extent (km) |
| --- | --- | --- | --- | --- |
| Rate of wild bee pollen deposition | Genung et al. 2022 | Landscape array of 3 plant species | Each plant species present at 25 sites | 35 |
| Fish biomass | Lefcheck et al. 2021 | 32 globally distributed ecoregions (16 temperate, 16 tropical) | 11-186 sites (median 59) | 17 - 4,677 |
| Above-ground carbon storage | Condit et al. 2000, Cavanaugh et al. 2014 | Tree species ID and estimated biomass at four globally distributed tropical forests | 50 1-Ha subplots from the 50-Ha BCI census; sets of six 1-Ha plots in three tropical regions. | 1 (Condit et al.); 32-681 (Cavanaugh et al.) |

### Which value of ℓ produces the strongest biodiversity-ecosystem function correlations?

We computed Hill diversity as a function of species relative abundances, *p_1, p_2,…, p_S*, and a scaling factor, ℓ, using the formula

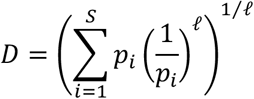

or its limit as ℓ approaches zero (exponential Shannon entropy) [18,50]. Historically, it has been more common to express Hill diversity with a scaling parameter “a” or “q”, equal to 1-ℓ [6,16,51]. We used the expression above (with the scaling parameter ℓ, instead), to highlight that this expression is a specific example of the more general weighted power mean [17,18]. This ℓ formulation’s biggest advantage over the q-formulation is that it clarifies the differences between weights (abundances) and the rarity scaling controlled by the parameter ℓ [18]. Furthermore, because the equations for richness, Hill Shannon, and Hill-Simpson diversities straddle ℓ = 0, this formulation may support the intuition that the spectrum of Hill diversity measures can extend in either direction (Figure 1), either further emphasizing common species (as ℓ gets increasingly negative) or further emphasizing rare ones (as ℓ gets increasingly positive).

We used observed species abundances to calculate species diversities at each site as the Hill diversity along a wide range of ℓ values (from −10 to +10 at intervals of 0.05) (Figure S1). We calculated total function as the sum of species’ functions at each site. We computed the correlation between the natural logarithm of each diversity and the natural logarithm of total function, across all sites in the community dataset (hereafter, the “BEF correlation”). We focused on the logarithms of function, and later, abundance variables because log(total function) = log(abundance) + log(per capita function), and we also log-transformed diversity because we anticipated that across the wide array of ecosystems considered, multiplicative, rather than additive, differences in diversity would be most comparable. To identify the ℓ value that produced the strongest BEF correlation in each community dataset, we plotted the correlation against the scaling factor ℓ. We identified the single ℓ value with the largest absolute correlation (i.e., largest R-squared for the relationship between log(diversity) and log(function)).

### How do biodiversity-ecosystem function correlations change in sign and strength over a wide range of Hill diversities?

To determine not only which ℓ value produced the strongest BEF correlation across community datasets, but also to see how adjusting the Hill diversity scaling parameter affects BEF relationships more comprehensively, we plotted the BEF correlation against the Hill diversity scaling factor ℓ for each community dataset. We examined curves to identify patterns in the sign and strength of the BEF correlation along the spectrum of emphasis on common and rare species.

### What is the role of absolute abundance in shaping BEF correlations?

To begin to separate effects of total and relative abundance on BEF correlations across the Hill diversity spectrum, we looked separately at the relationships between diversity and two complementary components of total function, namely total abundance and mean per-capita function. We used the same graphical approach we described above, regarding the sign and strength of the BEF correlation across the Hill diversity spectrum. For each community dataset, we found the correlation between the natural logarithm of Hill diversity at each site and either the natural logarithm of total abundance at each site, or the natural logarithm of mean per-capita function at each site, and plotted these correlations against the Hill diversity scaling parameter ℓ. Although on the logarithmic scale, abundance and mean per-capita function combine additively to create total function, the BEF correlation does not additively decompose into abundance by biodiversity and per-capita function by biodiversity correlations, as there is also covariance between abundance and per-capita function. Acknowledging this caveat, we suggest that by partitioning total function into additive components and examining how each of these relates to biodiversity gradients across the Hill spectrum, we can better characterize the role of total abundance in generating patterns in the BEF correlation itself.

## Results

### Which value of ℓ produces the strongest biodiversity-ecosystem function correlations?

For most datasets, the strongest biodiversity-ecosystem correlations were located at or just above richness (ℓ = 1), with a mode at ℓ = 1.5 (Figure 2). A substantial minority (11 of 39 datasets) had strongest BEF correlations at values of ℓ > 5, including a peak at ℓ = 10, the largest value of ℓ we considered. There were a few outliers: Two tree carbon storage datasets had their strongest BEF correlations near inverse Simpson (ℓ = −1) and exponential Shannon (ℓ = 0) diversities, and a single fish dataset had a strongest BEF correlation at ℓ = −10, the smallest value of ℓ we considered (Figure 2).

**Figure 2.**
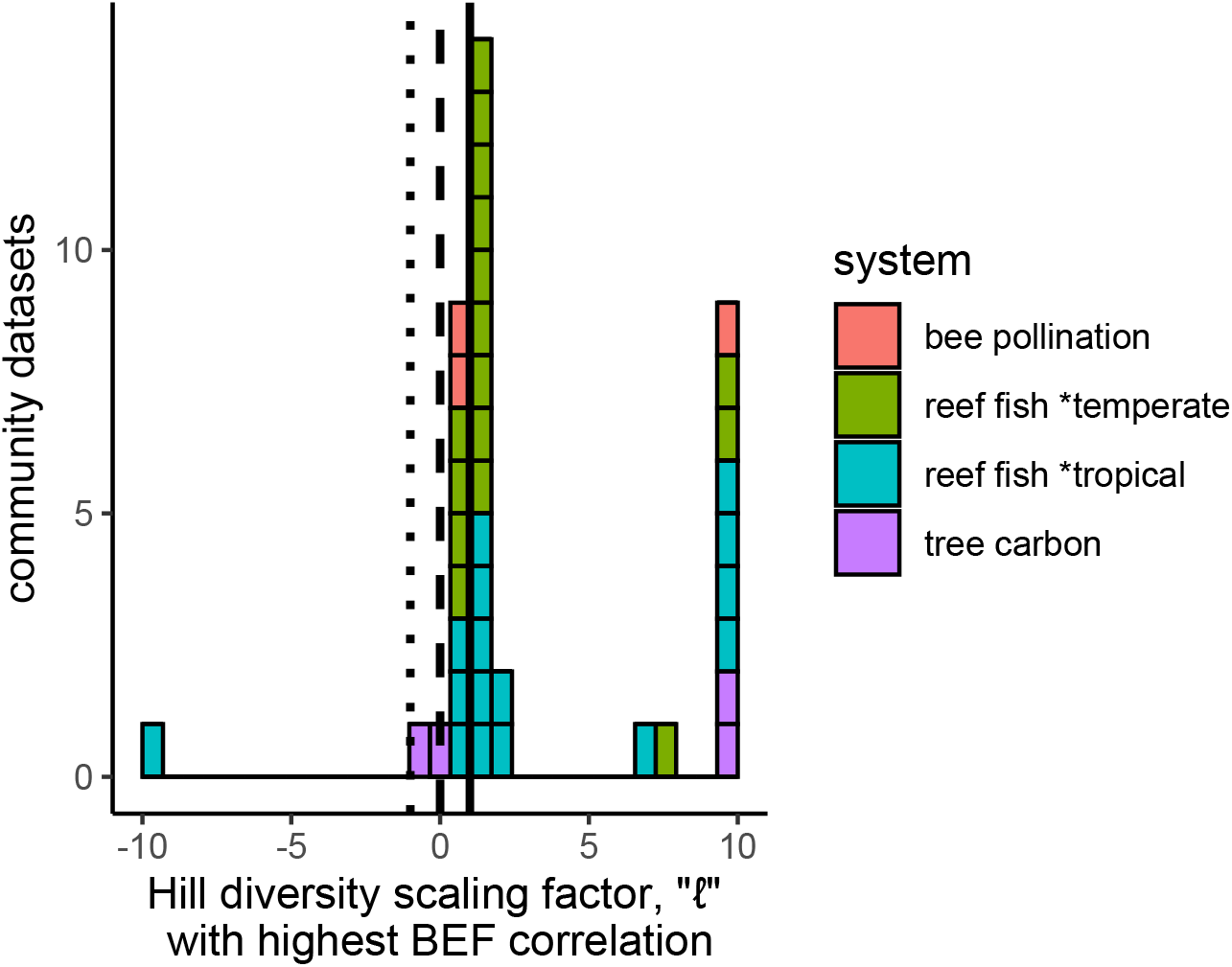
Across 39 observed biodiversity-ecosystem function (BEF) correlations calculated using a wide range of diversity scaling factors, the BEF relationships with the highest r^2^ were typically found using diversities near richness (vertical solid line; modal *ℓ* = 1.5). The highest BEF r**^2^** value for a community dataset was rarely found using diversities that emphasize the relative abundance of common species, including exponentiated Shannon (dashed line) and inverse Simpson (dotted line). Correlations were calculated between log(diversity) and log(ecosystem) function at a site (total above-ground carbon in tropical forest plots, rate of pollen grain deposition by wild bees, or total biomass of reef fish encountered in fixed-effort dive surveys in temperate and tropical regions).

### How do biodiversity-ecosystem function correlations change in sign and strength over a wide range of Hill diversities?

Across all ecosystem functions, we found common patterns in the relationship between the BEF correlation and the Hill diversity scaling parameter, ℓ. When ℓ was < 1, the diversity-function correlation was typically weak and could be positive or negative (Figure 3). Near ℓ = 1, the diversity function correlation rapidly increased, although a substantial minority of community datasets first showed a sharp negative turn in the relationship near Hill-Simpson and Hill-Shannon diversities (Figure 3 b-d). Across all the datasets we considered, the mean correlation between log(diversity) and log(total function) was not significantly different from zero for either Hill-Simpson or Hill-Shannon diversity (p>0.28 for two-sided Student’s t-test, with no correction for multiple tests). At richness (ℓ = 1), almost all datasets showed positive diversity-function correlations, with the mean R^2^ = 0.381. For most datasets, the strongest correlations were located near richness, with a mode near ℓ = 1.5, where the mean R^2^ was 0.445, after which the diversity-function correlation slowly declined as ℓ values continued to increase (Figure 3). A substantial minority of datasets showed continually stronger relationships as ℓ increased (some profiles in Figure 3 b, c), leading to highest R^2^ values at or near the maximum ℓ we considered (ℓ = 10).

**Figure 3.**
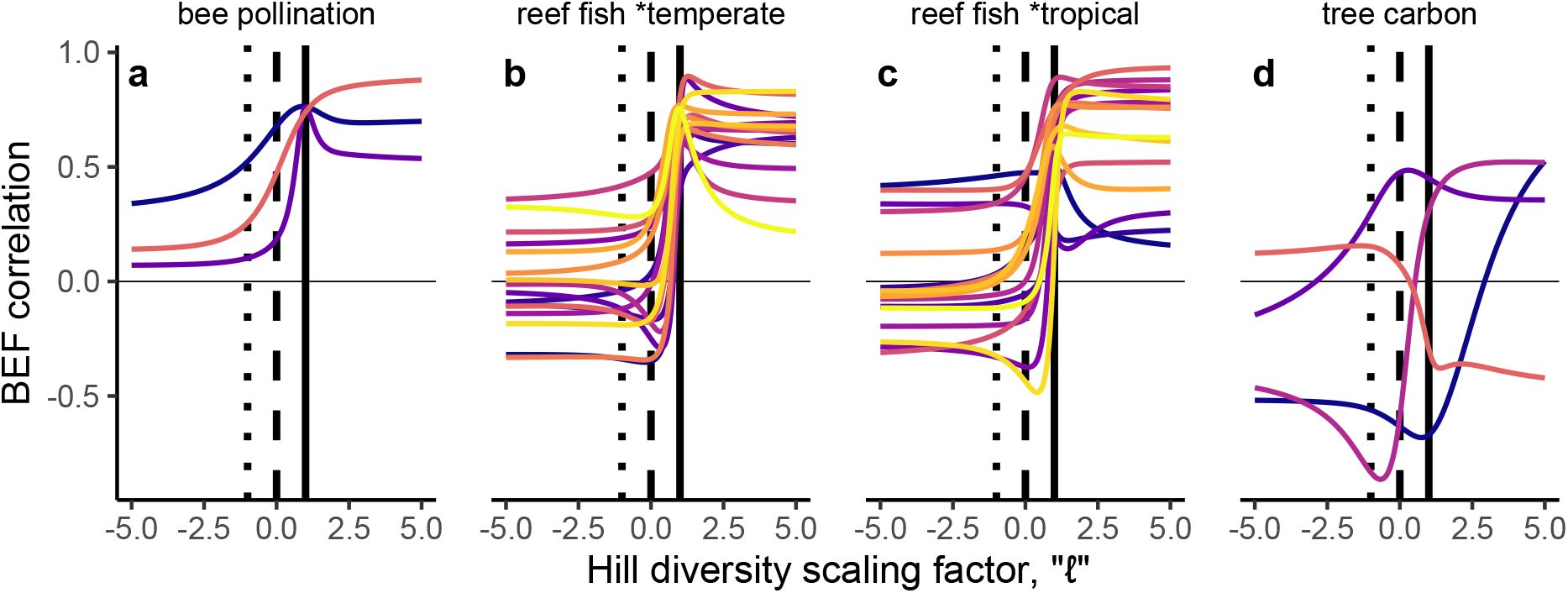
Biodiversity-ecosystem function (BEF) correlations across observed communities in a study system vary in magnitude and direction, depending on which scaling factor “ℓ” is used for calculating species’ diversity. Ecosystem services considered here are (a) rate of pollen grain deposition on one of three flower species by wild bees; total biomass of reef fish encountered in fixed-effort dive surveys in (b) temperate and (b) tropical global regions; and (d) total above-ground carbon in tropical forest plots. Correlations are between logged total function at each site, and logged species diversity at a range of ℓ values (at 0.05 intervals) emphasizing the relative abundance of common species’ (negative ℓ values) or rare species (positive ℓ values). Vertical lines identify correlations at commonly used diversities: inverse Simpson (dotted), exponential Shannon (dashed), and richness (solid). Colors visually distinguish different community datasets.

This study was not designed to contrast trends between ecosystem functions, but it is important to note that the relationship between the BEF correlation, and the emphasis the diversity metric puts on rare vs. common species (i.e., the value of ℓ), did not appear uniform across systems. For the three bee community datasets, total pollen deposition and bee diversity were positively correlated at every value of ℓ. Correlation strength peaked at richness (ℓ = 1) or just beyond (ℓ = 2), but remained relatively strong across all higher values of ℓ (Figure 3a). For the 32 reef fish community datasets, total fish biomass and fish diversity tended to be weakly and often negatively correlated at low ℓ values. Correlation strength tended to peak either slightly above richness at ℓ = 1.5, or grow with ℓ for an observed peak near the maximum value considered (ℓ = 10) (Figure 2, Figure 3 b, c). When considering either very high or very low ℓ values, note that at either end of the Hill number spectrum, diversities rapidly converged towards their maximum or minimum asymptote. Thus, large changes in the BEF correlation rarely occurred outside a fairly narrow range between ℓ = −2 and 2. Finally, the four tropical tree community datasets showed generally weak correlations. In two tree datasets, BEF correlation strength peaked at intermediate ℓ values where the BEF correlation was strongly negative (Figure 3 d). In another (Barro Colorado Island) diversity-function correlation was negative even at high ℓ values (Figure 3 d, orange line), but modestly positive for negative ℓ values.

### What is the role of absolute abundance in shaping BEF correlations?

As expected, the relationship between diversity and abundance was mostly similar to the relationship between diversity and function, as total abundance underlies function in our datasets. This can be seen in the similar shape of the curves showing the correlation between log(diversity) and either log(function) (Figure 3) or log(abundance) (Figure 4 a-d), as the sign and strength of correlation typically moved in similar ways across the ℓ spectrum. In almost all cases, the correlation between log(diversity) and log(abundance) was very strong (and in many cases approached unity), for large, positive values of the Hill diversity scaling parameter ℓ. As previously remarked, this result is a mathematical inevitability when datasets contain very rare species/singletons. Additionally, across datasets, we found that the rise towards the high correlation observed for large ℓ values typically occurred in the range of ℓ values typically considered by ecologists (−1 to 1), likely reflecting biological and sampling linkages between abundance and diversity; the correlation frequently saturated once ℓ was greater than two. While for some community datasets, diversity was largely independent of abundance for negative ℓ values, we also saw community datasets in which log(abundance) and log(diversity) had modest to strong negative correlation across negative ℓ values. Because Hill diversities typically change little with ℓ below around −2 [6], this result implies that in these systems, total abundance and the degree of dominance were positively linked [52].

**Figure 4.**
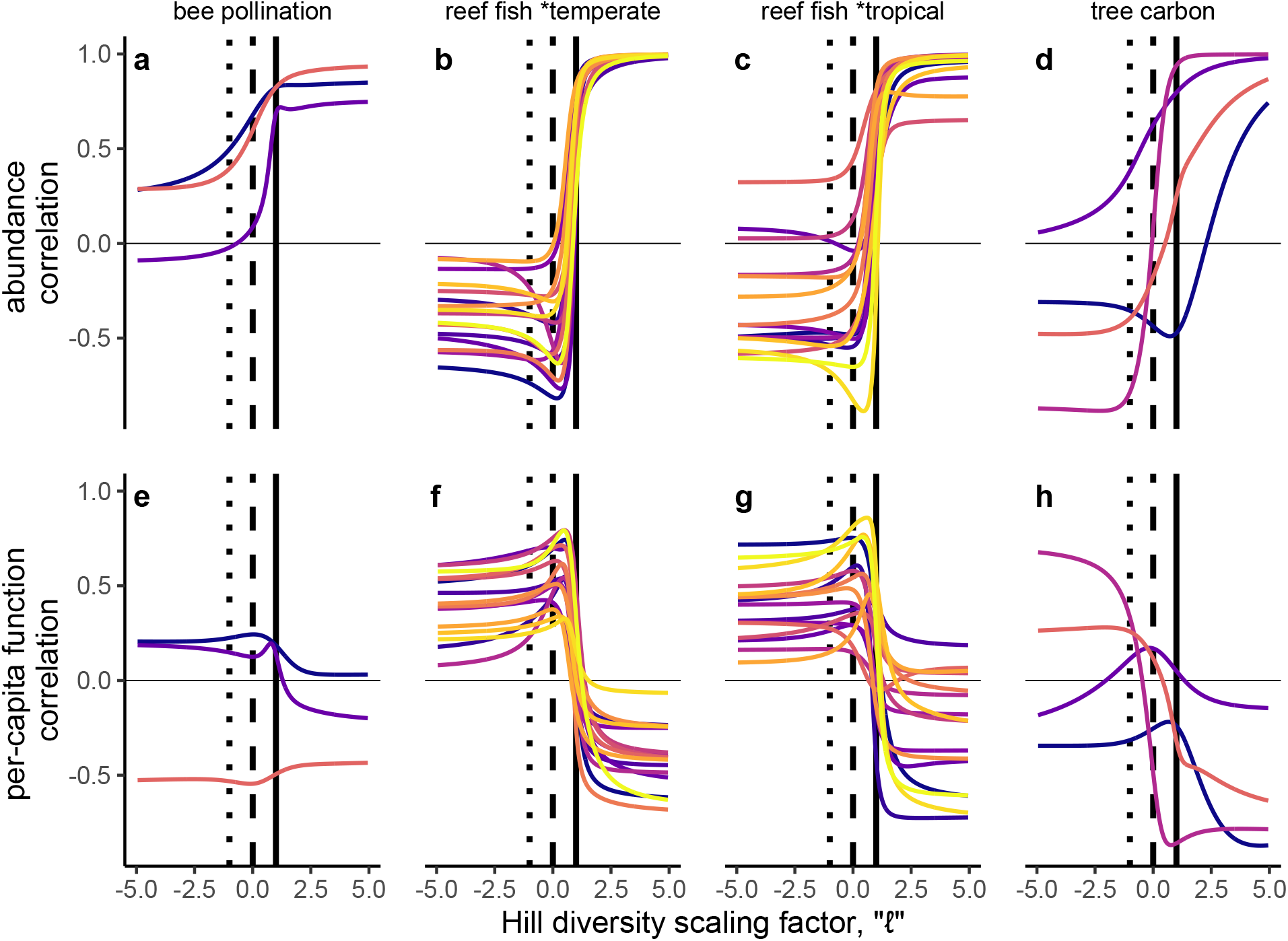
Biodiversity-ecosystem function (BEF) correlation in observational data (Figure 3) can be explained by correlation between total abundance and diversity (first row), correlation between mean per-capita function and diversity (second row), and interactions between these two factors (intractable, not shown). With a few exceptions, the abundance-diversity correlations roughly match the BEF correlation across the range of ℓ values used to calculate species diversities, while per-capita function-diversity correlations show countervailing trends in magnitude and direction. Compare (a, e) wild bee pollination, reef fish biomass in (b, f) temperate and (c, g) tropical regions, and (d, h) tropical forest above-ground biomass with corresponding panels in Figure 3. Vertical lines identify correlations at commonly used diversities: inverse Simpson (dotted), exponential Shannon (dashed), and richness (solid). Colors visually distinguish different community datasets.

While the curves in Figures 3 a-d and 4 a-d show strong resemblance, for some community datasets the BEF and diversity-abundance relationships diverged, implying those BEF relationships resulted from processes other than abundance. For example, correlations between abundance and diversity were always strongly positive for the reef fish data for large, positive ℓ values, but in some reef fish community datasets, correlations between diversity and function were only weakly positive at higher ℓ values (Figure 4 b, c). Such divergences between the diversity-function and diversity-abundance curves could be due to strong and/or countervailing relationships between mean per-capita function and diversity, which also showed some overall patterns across community datasets (Figure 4 e-h). In general, Hill diversities with negative ℓ values were positively related to per-capita function, suggestive of a positive relationship between evenness and mean per-capita function. This pattern was not ubiquitous, however, with notable exceptions in both tree and bee community datasets (Figure 4 e, h). We found that the correlation between diversity and mean per-capita function often exhibited a positive peak at intermediate ℓ values, a pattern particularly pronounced in the reef fish community datasets (Figure 4 f, g). Finally, there was a tendency towards a negative correlation between log(diversity) and log(mean per-capita function) for larger, positive values of ℓ (when diversity becomes largely synonymous with abundance), though the strength of this relationship was variable.

## Discussion

Diversities near richness (1 < ℓ < 2) often had strong positive correlations with ecosystem function (Figure 2), echoing a large body of research emphasizing the importance of richness in BEF relationships [3]. However, this was not a foregone conclusion, because most empirical BEF work does not consider other portions of the diversity spectrum [8], and because much of the theoretical and empirical work is grounded in closed communities where richness has a clear interpretation. We were surprised by the consistent performance of richness, because of three concerns we described in the introduction: first, observed richness is not a robust biodiversity measure in observational data; second, Hill diversities with high ℓ should better explain function in systems with highly variable total abundance; third, ℓ values that emphasize the probability of interspecific encounter (−1 ≤ ℓ < 1) might better explain function if species interactions are very important. The disconnect between observed and true richness is not, practically speaking, a resolvable problem and thus we cannot evaluate how much this first issue is affecting our results [25]. In the following paragraphs, we explore the latter two points, namely: how do Hill diversities near richness outperform Hill diversities with higher ℓ that better reflect abundance, and why we might have found such low explanatory power for Hill diversities that should capture the effects of (potentially positive) species interactions. Although all the information about relative abundances is contained in any continuous interval along the Hill diversity spectrum, at different values of ℓ, different aspects of the abundance distribution are emphasized. To better ground our discussion, we use an admittedly imprecise simplification, and refer to ℓ values as falling within the “inverse dominance range” (ℓ < 1), the “evenness emphasis range” (−1 ≤ ℓ < 1), the “rare emphasis range” (1 ≤ ℓ < 2, justification follows), or the “abundance emphasis range” (ℓ > 2).

As ℓ values increase, empirical diversities values are increasingly dependent on total abundance, since the abundance of the rarest species is typically at the lower bound set by the detection threshold (for count data, one). Hill diversities in the “rare emphasis range,” like richness itself, are affected by abundance as well as diversity *per se*. This is reasonably viewed as a sampling nuisance [6,24]. But viewed in another light, measures that combine abundance, relative abundance, and richness components could predict function well because, owing partly to their sampling properties, they describe salient biodiversity gradients. In fact, our results showed that ℓ values in the rare emphasis range that, compared to richness, are relatively more sensitive to abundance tended to better explain function. Rare-emphasis Hill diversities increase with richness, abundance, and often also dominance. As a result, they might vary with the importance of sampling effects whereby higher-functioning (likely, highly abundant) species are more likely to occur in species-rich assemblages (a kind of selection effect) [34,37,53]. We found that rare-emphasis Hill diversities tended to explain function best, with 1 < ℓ < 2 performing better than richness, ℓ = 1, but because they still reflect compositional heterogeneity, also better than abundance alone (Figures 2–3).

Hill diversities in the evenness emphasis range (−1 ≤ ℓ < 1) should capture the effects of species interactions by emphasizing the probability of interspecific encounter, but these diversities explained function poorly. This was partly unexpected because in several real-world systems, function increases with evenness [54–57], which increases Hill diversity for ℓ < 1 [58–60]. Additionally, in the evenness emphasis range, sample Hill diversities have relatively good statistical properties as estimators of true diversity, and asymptotic estimators [24] can further improve the situation, largely avoiding the robustness issues we highlight with species richness. Instead, the observed weak explanatory power of Hill diversities with ℓ values in the evenness emphasis range is because functions analyzed here are the product of two components, abundance and per-capita function, which each showed different responses to increasing ℓ. Abundance-diversity relationships often followed function-diversity correlations (compare Figure 3 with the top row of Figure 4). In other words, across the ℓ spectrum, Hill diversities had nearly the same relationship with abundance and with function, underlining the necessity of accounting for the role of total abundance in BEF research [61]. However, Hill diversities in the evenness emphasis range deviated from this pattern, instead exhibiting often strong, countervailing relationships with abundance and per-capita function (Figure 4). This fits with previous BEF literature, which anticipates a variety of mechanisms linking evenness and ecosystem function, without a clear prediction for when the net result is positive or negative [62–65].

The relationship between diversity and per-capita function differed from the diversity-abundance relationship, with correlation coefficients for per-capita function generally decreasing with increases in ℓ, but often showing a positive peak in the evenness emphasis range (Figure 4). Positive species interactions, including those that increase per-capita function, are expected to explain total function [53,66,67]. Our results partly support these expectations, as Hill diversities in the evenness emphasis range, which should track the probability of interspecific encounter, were positively associated with per-capita function, even as they tended to be negatively associated with abundance. As we increased ℓ, the correlation between Hill diversity and per-capita function disappeared near richness (ℓ = 1), also pushing against expectations that richness best captures function-relevant biodiversity gradients. In the rare emphasis and abundance emphasis ranges, we typically found a negative correlation between Hill diversity and per-capita function. Recent work highlighting the functional contribution of rare species to ecosystem function led us to suspect the opposite might occur [43,68–70]. The observed negative correlation likely reflects spatial constraints and/or fundamental tradeoffs between having many, smaller-bodied individuals versus fewer larger ones [71,72]. This scenario is particularly easy to imagine for trees crowding in fixed-area plots, which physically and energetically prohibit arbitrarily large numbers of the largest trees. Similar energetic and spatial constraints limit the number of very large fish that might be seen in a single dive. Thus, we suspect that one reason we saw a decline in the correlation between mean per-capita function and diversity with increasing ℓ in the fish and tree datasets is decreases in per-capita function due to crowding.

Even as Hill diversities in the rare emphasis range most often explained total function best, Hill diversities with ℓ values in the abundance emphasis range also explained function well, and should not be discounted. Hill diversities in the abundance emphasis range were the *best* predictor of function in a substantial minority of datasets (Fig. 2), and for nearly all datasets were *strong* predictors of function (Fig. 3, far right of x-axes). This was expected because of a general link between higher abundance and higher function [41,61,73–76]. Even as Hill diversities in the “abundance emphasis” range related strongly to function, we also note that abundance (like evenness) can relate to Hill diversity across the full spectrum of ℓ values. For example, if high-abundance sites tend to be dominated by many individuals of one or a few species [52], Hill diversities that emphasize inverse dominance will decrease with abundance. Thus, we should not expect that strong effects of abundance on function are captured exclusively at high values of ℓ.

The explanatory power of Hill diversity changed nonlinearly with increases in ℓ, as multiple facets of community structure (e.g., richness, abundance, evenness) affect function simultaneously. If we had found a monotonic strengthening of BEF relationships with increasing ℓ, we would argue that Hill “diversities” with large positive scaling parameters were simply abundance metrics masquerading as measures of diversity. Instead, we found, across a variety of regions, taxa, and ecosystem functions, intermediate, positive ℓ values in the “rare emphasis” range tended to produce the strongest BEF relationships (Fig. 3). All Hill diversities with positive ℓ values (including richness) tend to increase with both abundance and richness, which we argue can be a useful property, especially for BEF research. Because the goal of summarizing species’ abundances with diversity metrics is to distill complex, multivariate information [17], this claim is not radical. In fact, Hill diversities that emphasize rare species more than richness does can reflect intuitive notions of diversity, which include both high density and high compositional variation [77]. Our study points to the need for further theoretical work to explicate the meaning of these seldom-used Hill diversities in the rare emphasis range, and their linkages to ecosystem function.

By considering Hill diversities over a wide range of ℓ, we place ourselves at odds with the convention that Hill diversities should be considered only when ℓ ≤ 1 [6,16,17,51,78]. The most compelling argument for that restricted range of scaling parameters is presented by Patil and Taillie, who argued that diversity should not decrease when abundance is shifted from more to less abundant species, including to species with zero abundance, a variation on Dalton’s “principle of transfers” [17,79]. This diversity property does not hold for Hill diversity when ℓ >1, which has species richness as its minimum, occurring in the perfectly even community, and increases (given richness and abundance) as some species get progressively rarer. A more pragmatic argument comes from Chao et al., who noted that estimating the relative abundance of rare species is an increasing problem for diversity measures as ℓ increases; they therefore suggest using only more estimable Hill diversities with ℓ ≤1 [6]. However, theoretical work suggests that even richness (ℓ = 1) is poorly estimated [24,25], and by this logic should not be used either. Finally, and most generally, diversity measures have traditionally been considered separate from abundance/density measures (but see [80,81]), whereas with increasing ℓ values, observed diversity and observed abundance tend to be more strongly correlated (and in fact approach a correlation of one in our datasets). Despite these arguments, our results show that Hill diversities with ℓ > 1 are meaningful ecological diversity measures, at least in the sense that they convey more information about function than do more widely-used Hill numbers. Choosing to exclude these Hill diversities might be desirable for some conceptualizations of diversity, but we are opening the narrower question of which Hill diversities—with their variable emphasis on richness, abundance, evenness, and dominance—best explain ecosystem function. In this pursuit, allowing diversity metrics to highlight absolute abundance is valuable.

Our correlational analyses of observational data do not consider confounding variables, which may obscure links between biodiversity and ecosystem function, across the Hill diversity spectrum. Experimental and statistical approaches to better account for environmental drivers of diversity and/or function and more rigorously trace causal pathways (e.g., [38]) will be useful in validating and extending our findings [82]. One obvious effect of ignoring confounding variables is that the BEF correlations we found are likely to be low, as the confounding variables add noise that is not accounted for. Future work linking environmental gradients and other confounding variables to particular regions of the Hill diversity spectrum (e.g., regions emphasizing inverse dominance, evenness, rare species, or total abundance) may also clarify mechanisms underlying BEF relationships.

As global changes lead to shifting species abundances, ecologists must continue to describe and predict how these shifts impact ecosystems and the way they function. Yet, understanding the separate and combined roles of total and relative abundance in mediating ecosystem function remains a difficult challenge, in large part because total abundance is inextricably linked to diversity measures. It is mathematically linked for large, positive ℓ values. It is practically constrained by sampling effects for ℓ closer to 1 (i.e., near species richness). As ℓ becomes negative, Hill diversities may lose their dependence on total abundance [24]. However, in the majority of community datasets, we saw at least weak negative correlations between negative-ℓ Hill diversities and observed abundances, likely due to increasing dominance in more abundant systems [52]. Overall, this suggests that in observational contexts, simple partitioning of abundance and diversity effects may not be tractable, at least not in a satisfying manner [34,35]. Since no single-best diversity measure is likely to emerge for all BEF studies, we encourage researchers to be open-minded towards Hill diversities across a wide spectrum of ℓ values and their potential links to mechanisms underlying BEF relationships.

## Acknowledgements

We thank Rachael Winfree for key discussions that sparked this work, Jonathan Dushoff for providing insight on Hill numbers, and participants in the Hakai Synthesizing Biodiversity Seminar Series for useful feedback on preliminary results. Mary O’Connor and two anonymous reviewers provided helpful feedback on an earlier draft of this manuscript. We gratefully acknowledge all data collectors and data holders. Data for three tropical forests (Pasoh, Indonesia; Yasuní, Ecuador; and Volcán Barva, Costa Rica) were originally made available to us by the Tropical Ecology Assessment and Monitoring (TEAM) Network, a collaboration between Conservation International, the Missouri Botanical Garden, the Smithsonian Institution, and the Wildlife Conservation Society, and partially funded by these institutions, the Gordon and Betty Moore Foundation, and other donors.

---

**Figure S1.**
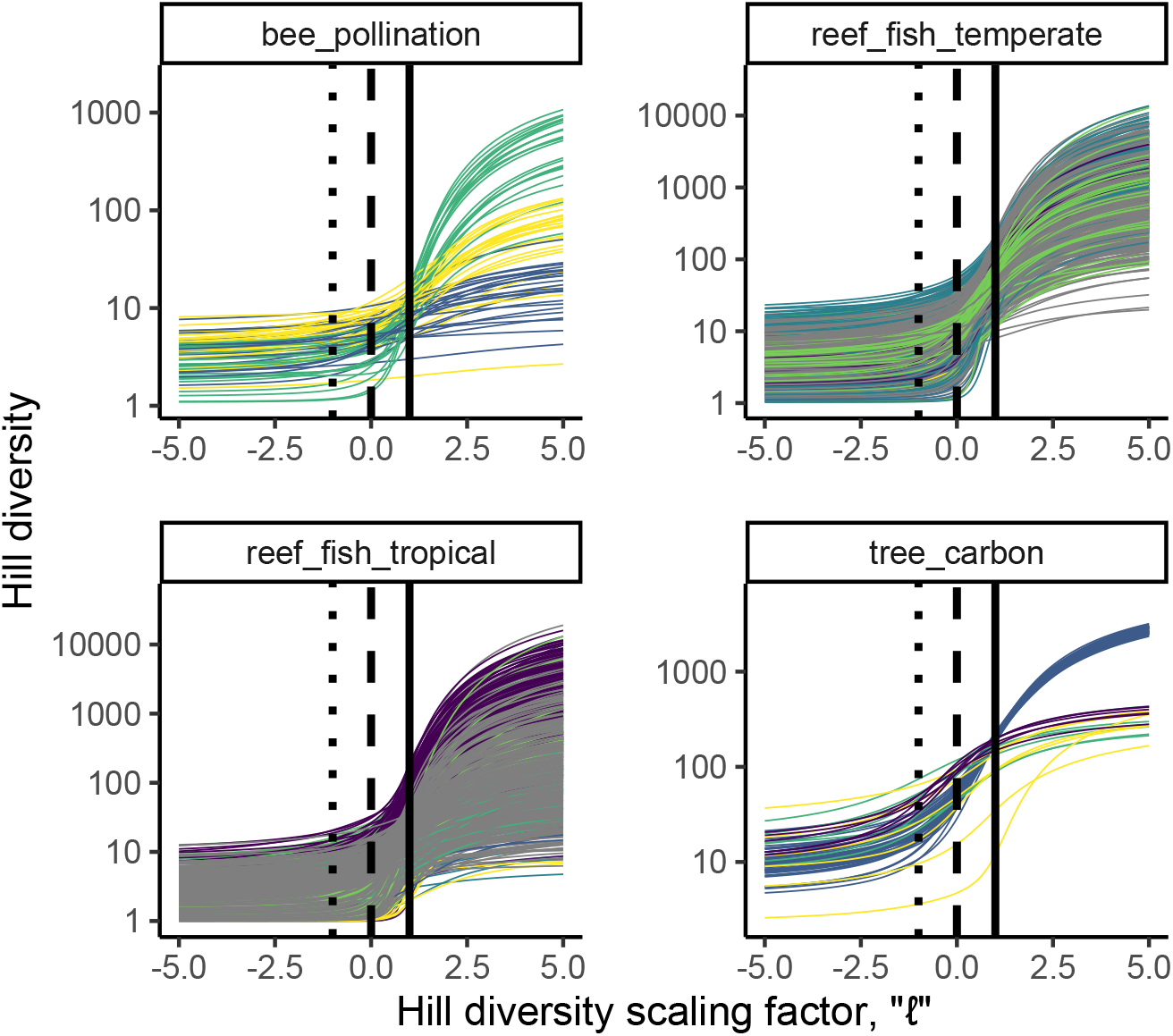
Community datasets (line color) differed in both the shapes of the diversity profiles (Hill diversity vs. ℓ) and the degree to which diversity profiles differed between sites. For example, the grey-blue tree_carbon sites were all 1-Ha subplots from the contiguous BCI 50-Ha forest plot, and diversity profiles were very similar between subplots; by contrast the yellow bee_pollination sites (Floral visitors of *Polemonium reptans*) had variable structure with wide variety in richness (ℓ = 1, vertical solid lines), inverse dominance (large negative ℓ), and abundance (large positive ℓ). Each curve is the diversity profile for a single site; colors indicate a community dataset (set of sites within a region at which a single function was measured).

